# CCR5 closes the temporal window for memory linking

**DOI:** 10.1101/2021.10.07.463602

**Authors:** Yang Shen, Miou Zhou, Denise Cai, Daniel Almeida Filho, Giselle Fernandes, Ying Cai, Nury Kim, Deanna Necula, Chengbin Zhou, Andy Liu, Xiaoman Kang, Masakazu Kamata, Ayal Lavi, Shan Huang, Tawnie Silva, Won Do Heo, Alcino J. Silva

## Abstract

Real world memories are formed in a particular context and are not acquired or recalled in isolation ^1-5^. Time is a key variable in the organization of memories, since events experienced close in time are more likely to be meaningfully associated, while those experienced with a longer interval are not^1-4^. How does the brain segregate events that are temporally distinct? Here, we report that a delayed (12-24h) increase in the expression of the C-C chemokine receptor type 5 (CCR5), an immune receptor well known as a co-receptor for HIV infection^6,7^, following the formation of a contextual memory, determines the duration of the temporal window for associating or linking that memory with subsequent memories. This delayed CCR5 expression in mouse dorsal CA1 (dCA1) neurons results in a decrease in neuronal excitability, which in turn negatively regulates neuronal memory allocation, thus reducing the overlap between dCA1 memory ensembles. Lowering this overlap affects the ability of one memory to trigger the recall of the other, thus closing the temporal window for memory linking. Remarkably, our findings also show that an age-related increase in CCL5/CCR5 expression leads to impairments in memory linking in aged mice, which could be reversed with a CCR5 knockout and an FDA approved drug that inhibits this receptor, a result with significant clinical implications. All together the findings reported here provide the first insights into the molecular and cellular mechanisms that shape the temporal window for memory linking.

---

Memory formation can be affected by previous experiences. For example, memories acquired close in time often become linked such that the retrieval of one increases the likelihood of retrieving the other (i.e., memory linking). Abnormal memory linking (e.g., improper relational memory), is involved in psychiatric disorders such as post-traumatic stress disorder and schizophrenia^8,9^. However, very little is known about the mechanisms that regulate the interactions amongst memories. Activation of CREB and subsequent increases in neuronal excitability are thought to open the temporal window for memory linking, so that a given neuronal ensemble involved in encoding one memory is more likely to participate in encoding a second memory acquired close in time^2,10-13^. Accordingly, the neuronal overlap between memory ensembles has been shown to be critical for memory linking^1-3^. In contrast, little is known about the mechanisms that segregate events that are temporally distinct. CCR5 has been extensively studied in the context of inflammatory responses and HIV infection^6,7^. However, comparatively little is known about its role in learning and memory. Both CCR5 and its ligand CCL5 are highly enriched in the CA1 region of the hippocampus^14,16^ and CCR5 is a negative regulator of CREB activation and neuronal excitability^15,17^. Here, we demonstrate that a gradual increase in the expression of CCL5/CCR5 following memory formation closes the temporal window for memory linking, thus segregating memories that are temporally distinct.

## CCR5 expression is enhanced after learning

To explore CCR5’s role in contextual memory linking, where the memory of one context is associated or linked to the memory of a second context^1^, we first tested whether the expression of CCR5 and its ligands change after contextual conditioning (Fig. 1a) in a brain region critical for this form of learning (i.e., dCA1). Compared to expression levels in mice that stayed in their home cage (HC), both *Ccr5* and *Ccl5* mRNA increased 12 hours (12h) after contextual conditioning (Fig. 1b, c), while there were no significant changes in the expression of other CCR5 ligands tested (*Ccl3, Ccl4* and *Ccl11*; Extended Data Fig. 1a-d). Next, we used *in situ* hybridization to determine the hippocampal cellular distribution of this learning-induced increase in *Ccr5* expression (Fig. 1d). Although in dCA1 of HC mice there were more *Ccr5*-expressing microglia than *Ccr5*-expressing neurons (Fig. 1e), there was a dramatic increase in *Ccr5*-expressing neurons, but not microglia, at 6h and 12h after contextual conditioning (Fig. 1f).

**Fig. 1.**
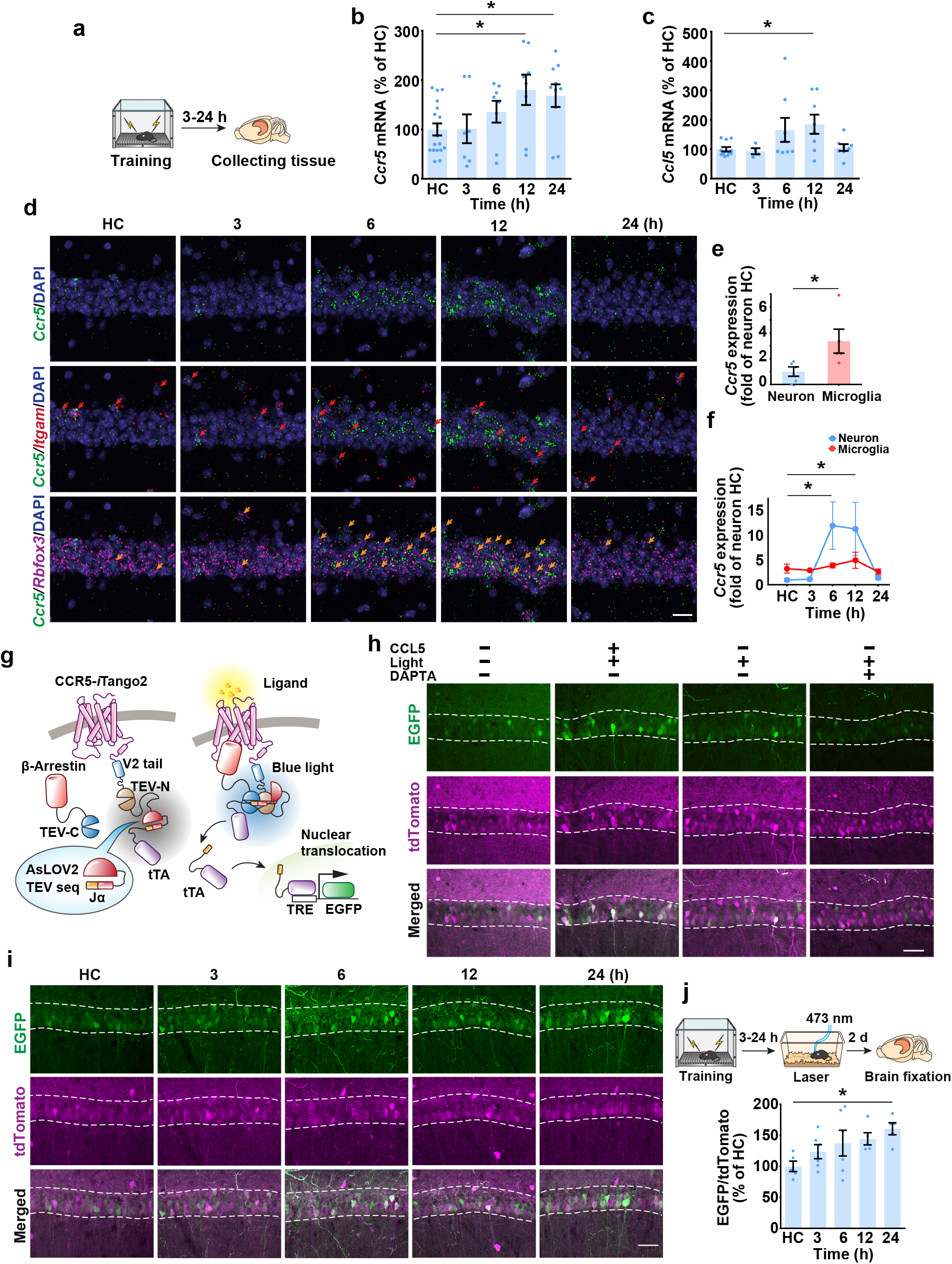
CCR5 expression and activation in the dorsal hippocampus after contextual fear conditioning. **a**-**c**, mRNA levels of *Ccr5* (**b**) and *Ccl5* (**c**) in mouse dCA1 at 3-24h after fear conditioning (**a**). Tissue (dCA1) from home cage (HC) mice was collected at the same time points (3-24h) and pooled together as the control HC group. Results were normalized to HC. (n=8-18 mice per group, \**P*<0.05, one-way ANOVA). **d**, Representative images of *Ccr5, Itgam* (microglia marker), and *Rbfox3* (neuronal marker) mRNA expression in dCA1 from naïve mice or mice 3-24h after fear conditioning. Scale bars, 20 μm. **e**, Number of *Ccr5*-expressing microglia and neurons in naïve mice (n=5 mice, \**P*<0.05, Student’s t-test) **f**, Number of *Ccr5*-expressing microglia and neurons 3-24h after fear conditioning (n=4-5 mice, \**P*<0.05, two-way ANOVA). **g**, Schematics for CCR5-*i*Tango2. **h**, Representative images of CCR5-*i*Tango2-expressing dCA1 neurons after treatment with CCL5, DAPTA (CCR5 antagonist) and light stimulation. Scale bar, 50 μm **i**, Representative images of CCR5-*i*Tango2 expressing dCA1 neurons after fear conditioning. Scale bar, 50 μm. **j**, Quantification of EGFP expression (intensity normalized to tdTomato which is tagged to β-Arrestin through P2A, reflecting expression of the *i*Tango system. n=5-6 mice, \**P*<0.05, one-way ANOVA). All results shown as mean ± s.e.m.

In addition to CCR5 expression, we also measured neuronal CCR5 activity after learning with the *i*Tango2 approach^18^ (Fig. 1g). The light- and ligand-gated gene expression system we constructed (CCR5-*i*Tango2) enables cellular expression of a reporter gene (i.e. EGFP) only in the presence of both CCR5 ligand(s) and blue-light exposure (detailed information in Methods). When tested in either HEK293 cells (Extended Data Fig. 2), in dCA1 (Fig. 2h), or in the dentate gyrus (Extended Data Fig. 3), CCR5-*i*Tango2 showed a significant increase in EGFP expression only when both light and ligand (CCL5) were present, demonstrating that CCR5-*i*Tango2 is capable of monitoring CCL5 dependent CCR5 activation. Therefore, CCR5-*i*Tango2 viruses were injected into mouse dCA1, and 3-weeks later mice were trained with contextual fear conditioning. Compared to HC controls, neuronal CCR5 activity in trained mice showed a gradual increase after conditioning (Fig. 1i, j), a result consistent with the delayed expression patterns of CCR5 and CCL5 presented above (Fig. 1b, c and f). Overall, our results demonstrated that after contextual learning there was a delayed (12-24h) increase in CCL5/CCR5 signaling in dCA1 neurons.

**Fig. 2.**
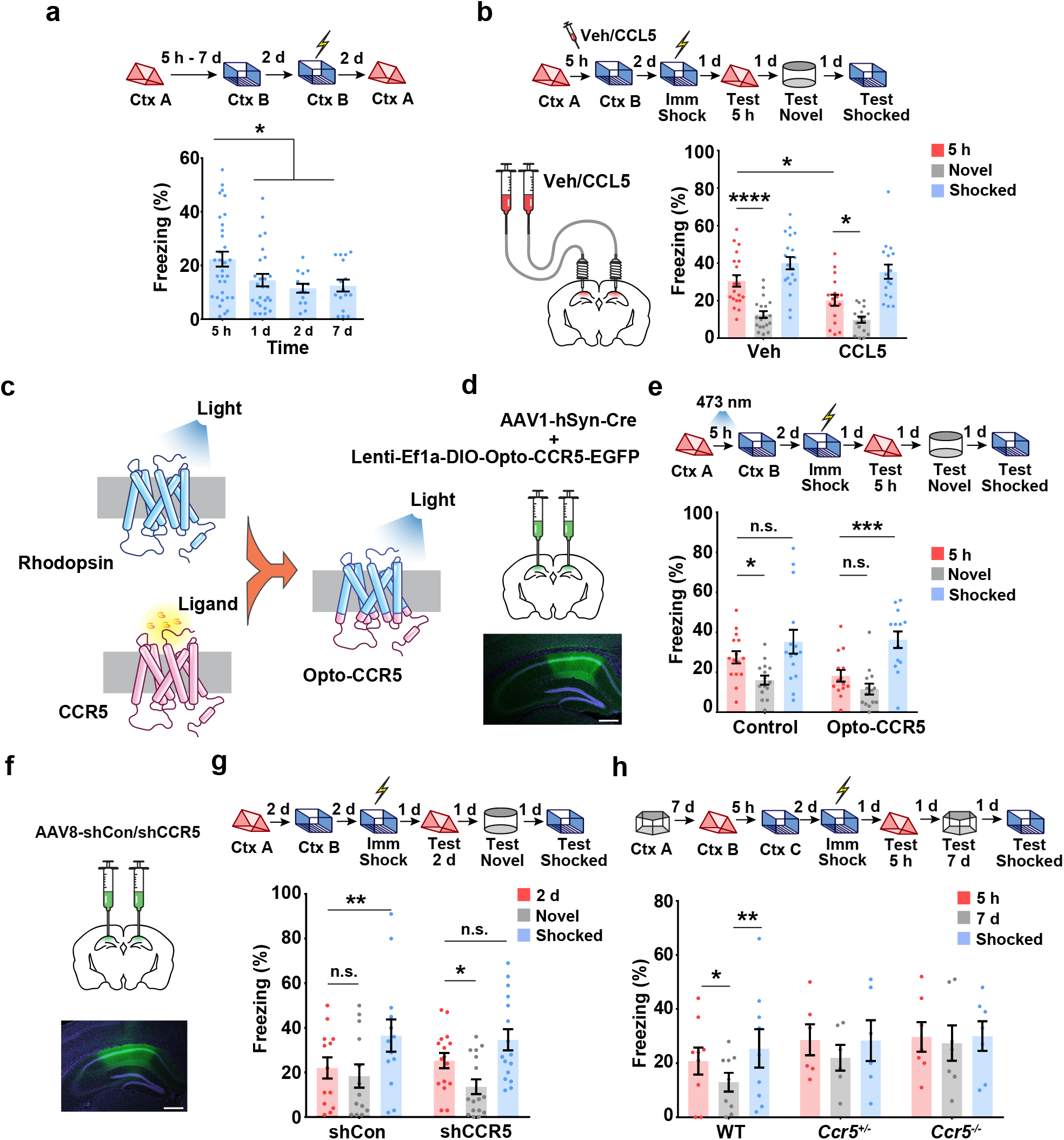
CCR5 regulates the temporal window of memory linking. **a**, Characterization of the temporal window for contextual memory linking (Ctx A, Context A; Ctx B, Context B; 5h n=32, 1d n=26, 2d n=14, 7d n=16, \**P*<0.05, one-way ANOVA). **b**, CCL5 infusion in dCA1 impaired contextual memory linking with a 5h interval between memories (Veh n=20, CCL5 n=17; \**P*<0.05, \*\*\*\**P* <0.0001, two-way repeated measures ANOVA). **c**, Schematics of the Opto-CCR5 construct. **d**, Schematics of viral constructs injection. Scale bar, 500 μm. **e**, Optogenetic activation of neuronal CCR5 impaired 5h contextual memory linking (Control n=15, Opto-CCR5 n=14; \**P*<0.05, \*\*\**P*<0.001, two-way repeated measures ANOVA). **f**, Schematics of AAV8-shCCR5 intrahippocampal injection. Scale bar, 500 μm. **g**, *Ccr5* knockdown in dCA1 neurons extended the temporal window of contextual memory linking (shRNA-Cont n=14, shRNA-CCR5 n=16, \**P*<0.05, \*\**P*<0.01, two-way repeated measures ANOVA). **h**, *Ccr5* knockout extended the temporal window of contextual memory linking (WT n=9, *Ccr5*^+/-^ n=6, *Ccr5*^-/-^ n=7, \**P* < 0.05, \*\**P* < 0.01, two-way repeated measures ANOVA). All results shown as mean ± s.e.m.

## CCR5 regulates contextual memory linking

To determine whether CCR5 modulates the temporal window for contextual memory linking^1^, we first exposed the mice to one context (context A) and either 5h, 1, 2 or 7 days later we exposed the mice to a second context (context B) (Fig. 2a). Two days later, the mice were given an immediate shock in context B, and then contextual memory linking was tested 2d later in context A. During the memory linking test, the 5h group showed higher freezing (i.e. higher linking) than the 1d, 2d or the 7d groups. This result shows that contextual memory linking decreases significantly between 5h and 24h, indicating a time course parallel to the increase in CCR5 signaling after learning (Fig. 1). Therefore, we subsequently investigated whether increasing or inhibiting CCR5 signaling affected the temporal window for contextual memory linking.

We first enhanced CCR5 activity by infusing CCL5 into dCA1 4h after mice were exposed to context A, a time point that preceded the expected endogenous CCR5 signaling increase. During the contextual memory linking test, compared to the control group, the CCL5 group showed significantly lower freezing in context A that the mice had explored 5h before context B (Fig. 2b), indicating that increasing CCR5 activity led to an attenuation of contextual memory linking. We then tested whether contextual memory linking could be regulated specifically by direct manipulation of neuronal CCR5 activity with a genetically encoded optical tool (Opto-CCR5) with high spatiotemporal precision^19^ (Fig. 2c; detailed information in Methods). Consistent with CCR5 activation^20-23^, light stimulation of Opto-CCR5 caused both a significant enhancement of intracellular Ca^2+^ and phosphorylation of Erk1/2 (Extended Data Fig. 4). To ensure specific neuronal expression, AAV1-hSyn-Cre was co-injected with Lenti-EF1a-DIO-Opto-CCR5 (or EGFP control virus) into dCA1 (Fig. 2d and Extended Data Fig. 5). During the contextual memory linking test, only the control group, but not the Opto-CCR5 group, showed evidence of memory linking (i.e., higher freezing in context A, that the mice experienced 5h before context B, compared to a novel context; Fig. 2e), confirming that increasing neuronal CCR5 activity specifically after exposure to context A is sufficient to impair contextual memory linking without impairing memory for context B.

To examine whether attenuating CCR5 signaling could extend the window for contextual memory linking, AAV8 containing shRNA-Control or shRNA-CCR5 was injected into dCA1 (Fig. 2f). Three weeks later, mice were pre-exposed to context A and then context B with a 2d interval. As expected, during testing, the control group did not show memory linking (i.e., showed similar freezing in context A as in a novel context; Fig. 2g). In contrast, the shRNA-CCR5 group showed higher freezing in context A than in a novel context, and there was no difference in freezing between contexts A and B, demonstrating strong memory linking (Fig. 2g).

*Ccr5* knockout mice (*Ccr5*^-/-^ mice) were also tested for contextual memory linking. As expected, during the test in context A, the WT mice froze less when the interval between contexts was 7d versus 5h. In contrast, *Ccr5*^*-/-*^ mice showed similar freezing in context A when the intervals between context A and B were 5h or 7d. These freezing levels were also similar to those shown in the shocked context (context B; Fig. 2h), demonstrating strong memory linking in *Ccr5*^*-/-*^ mice with a time interval (i.e., 7d) when WT mice do not show memory linking. Thus, two very different manipulations that decreased the levels of CCR5 (shRNA and a knockout) extended the temporal window for memory linking. Altogether, our results showed that increasing or inhibiting CCR5 signaling impaired or extended (respectively) the temporal window for contextual memory linking, demonstrating a key role for CCR5 in setting the duration for the memory linking window.

## CCR5 modulates memory co-allocation

Next, we investigated how CCR5 regulates the temporal window for contextual memory linking. Previous results suggested that a temporary increase in neuronal excitability following learning^24,25^ biases the allocation of a subsequent memory to the neuronal ensemble encoding the initial memory^1^, and that this ensemble overlap was critical for memory linking^26^. Thus, we examined whether CCR5 modulated neuronal excitability and consequently ensemble overlap, since this could explain CCR5’s role in shaping the temporal window for memory linking. When treated with CCL5, CA1 neurons from acute hippocampal slices showed a decrease in current injection-induced firing rate (Fig. 3a, b), indicating an inhibition of neuronal excitability. This is a significant result, since neuronal excitability is critical for determining which specific neurons in a neural network will store a given memory (known as memory allocation)^13,27,28^. Importantly, decreases in excitability, caused by increases in CCR5 signaling following learning, could explain how this receptor closes the window for memory linking.

**Fig. 3.**
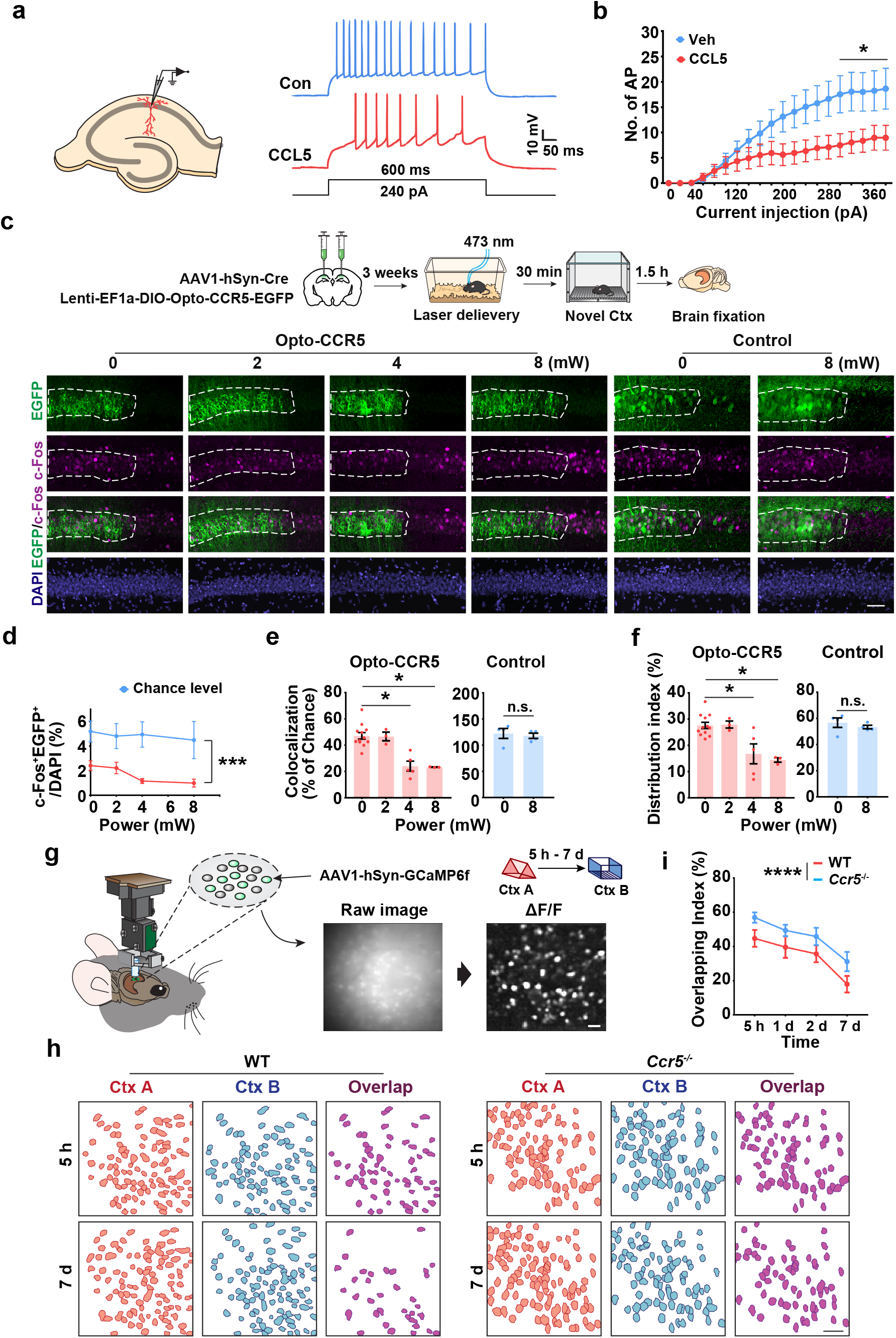
CCR5/CCL5 modulate neuronal excitability, c-Fos expression and the overlap of neural ensembles. **a**, Schematics of neuronal recordings and representative traces. **b**, CA1 neurons treated with CCL5 for 1h showed a significant decrease in firing rate (Control n=10 cells, CCL5 n=9 cells, \**P*<0.05, two-way ANOVA). **c**, Representative images of colocalization between c-Fos and Opto-CCR5-EGFP or EGFP control after light stimulation and novel context exposure. Scale bar, 50 μm. **d**, Percentage of c-Fos^+^EGFP^+^ cells at different power levels (0 mW n=13, 2 mW n=3, 4 mW n=5, 8 mW n=3, \*\*\**P*<0.001, two-way ANOVA). **e**, Percentage of colocalization between c-Fos^+^ cells and EGFP^+^ cells after normalization to chance level. Chance level = (c-Fos^+^/DAPI)*(EGFP^+^/DAPI)% (\**P*<0.05, one-way ANOVA). **f**, Quantification of c-Fos distribution in EGFP^+^ and non-EGFP cells in the Opto-CCR5-EGFP or EGFP control group. Distribution index = (c-Fos^+^EGFP^+^/EGFP^+^)/(c-Fos^+^EGFP^+^/EGFP^+^ + c-Fos^+^EGFP^-^/EGFP^-^)% (\**P*<0.05, one-way ANOVA). **g**, Schematics for miniscope setup and calcium signal identification. Images were collected from mice exploring different contexts separated by either 5h, 1d, 2d, or 7d. Scale bars, 50 μm. **h**, Neuronal overlap between different contexts. Scale bars, 50 μm. **i**, Overlapping index for WT and *Ccr5*^*-/-*^ mice (WT n=6, and *Ccr5*^*-/-*^ n=6; WT vs *Ccr5*^*-/-*^ mice, \*\*\*\**P*<0.0001, two-way ANOVA). All results shown as mean ± s.e.m.

To directly test whether increases in CCR5 activity could decrease memory allocation, Opto-CCR5-EGFP or EGFP control were expressed in mouse dCA1, and then subjected to blue light for 30 min (at different light power levels) before context exploration (Fig. 3c). Following light activation (4 and 8 mW) and contextual training, dCA1 neurons expressing Opto-CCR5 showed a significant reduction in the expression of learning-induced c-Fos, a widely used marker for neurons involved in memory^29^ (Fig. 3d-f), while the number of overall c-Fos^+^ cells were similar among groups (Extended Data, Fig. 6). This result is consistent with the hypothesis that CCR5 activation excludes neurons from memory ensembles. Additionally, light activation did not cause any changes in c-Fos expression in the GFP^+^ cells in the EGFP control group (Fig. 3e, f). Furthermore, when AAV8 containing shRNA-CCR5 was injected into dCA1, neurons with *Ccr5* knockdown had a higher probability of expressing c-Fos (i.e., being involved in memory; Extended Data, Fig. 7) compared with control neurons, a result that also supports the hypothesis that CCR5 activity modulates memory allocation in neuronal networks.

Altogether, the results presented suggest that the increase in CCR5 expression and signaling after learning prevents subsequent memories from being allocated to the neuronal ensemble encoding the initial memory, thus reducing the overlap between the two memory ensembles, and consequently attenuating memory linking. To further test this hypothesis, we recorded neuronal calcium activity (with GCaMP6f) in dCA1 with head mounted fluorescent microscopes (miniscopes^1^) while mice were exploring two different contexts separated by either 5h, 1d, 2d, or 7d. Then, we measured the overlap between the active neuronal populations recorded during the two contextual exposures in both WT and *Ccr5* knockout mice (Fig. 3g, h). Compared to WT mice, *Ccr5*^-/-^ mice revealed an overall significantly higher neural ensemble overlap (Fig. 3i; Extended Data, Fig. 8). Altogether these results strongly support the hypothesis that CCR5 modulates the temporal window for memory linking by regulating neuronal co-allocation and consequently the overlap between memory ensembles.

## CCR5 and aging-related linking deficits

CCR5 and CCL5 expression in peripheral immune cells increases with age^30,31^. Similar increases in aging neurons could contribute to age-related decreases in contextual memory linking^1^. To test this hypothesis, we measured hippocampal CCR5 and CCL5 expression in 16∼18-month-old mice (middle aged), an age in which mice still show intact contextual conditioning, but deficits in contextual memory linking^1^. Compared with young mice, middle-aged home cage mice had significantly enhanced *Ccl5* and *Ccr5* mRNA levels (Fig. 4a). Middle-aged mice also showed an increase in the transient *Ccl5* expression at 3h following contextual learning (Fig. 4b), which was earlier than young mice (6-12h after learning, Fig. 1c).

**Fig. 4.**
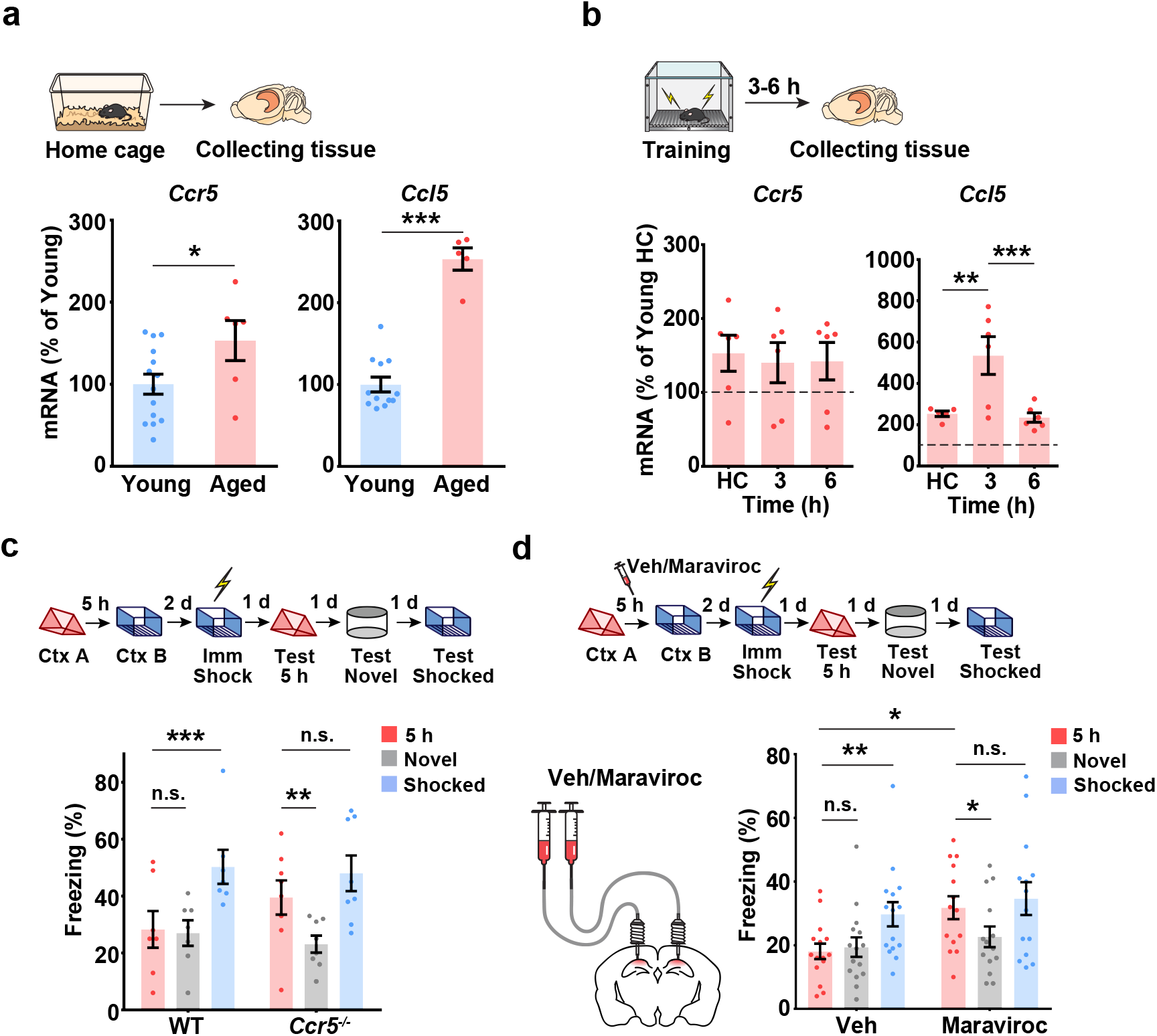
Enhanced CCL5/CCR5 signaling contributes to age-related memory linking deficits. **a**, Aged HC mice had higher *Ccr5* and *Ccl5* mRNA levels in dCA1 than young HC mice (*Ccr5*: young n=14, aged n=6, *Ccl5*: young n=12, aged n=5, \**P*<0.05, \*\*\**P*<0.001, Student’s t-test). **b**, *Ccr5* and *Ccl5* expression after fear conditioning in dCA1 of aged mice (*Ccr5*: n=6 for all groups, *Ccl5*: HC n=5, 3h n=6, 6h n=6, \*\**P*<0.01, \*\*\**P*<0.001, one-way ANOVA). **c**, *Ccr5* knockout rescued 5h memory linking deficits in aged mice (WT n=7, *Ccr5*^-/-^ n=8, \*\**P*<0.01, \*\*\**P*<0.001, two-way repeated measures ANOVA). **d**, Maraviroc, a CCR5 antagonist, rescued 5h memory linking deficits in aged mice (n=16 for each group, \**P*<0.05, \*\**P*<0.01, two-way repeated measures ANOVA). All results shown as mean ± s.e.m.

Although middle-aged mice showed deficits in contextual memory linking^1^, even when short intervals (i.e., 5h) were used (Fig. 4c), *Ccr5*^-/-^ mice showed clear evidence for memory linking tested with a 5h interval (i.e., higher freezing in contexts A and B than in a novel context; Fig. 4c). To test the effect of pharmacologically blocking CCR5 activity on contextual memory linking in middle-aged mice, maraviroc (an FDA approved CCR5 antagonist used for HIV treatment)^32^ was infused to dCA1 of these mice 1h before they were exposed to context B in a contextual memory linking experiment with a 5h interval. Unlike control mice, maraviroc-treated mice showed memory linking (Fig. 4d). Thus, blocking CCR5 with maraviroc ameliorates the memory linking deficits in middle-aged mice. Altogether these results support a role for CCR5 expression in closing the temporal window for memory linking as well as in age-related deficits in memory linking.

In summary, the findings reported here show that a delayed (12-24h) increase in the CCL5/CCR5 signaling in dCA1 neurons of a given memory ensemble closes the temporal window for contextual memory linking. CCR5 activation decreases neuronal excitability, thus negatively regulating memory allocation. This change in memory allocation decreases the overlap between dCA1 memory ensembles, and therefore impairs the ability of one memory to trigger the recall of the other, thus closing the temporal window for memory linking (Extended Data Fig. 9). Remarkably, our findings also show that an age-related increase in CCL5/CCR5 expression leads to impairments in contextual memory linking in middle-aged mice that could be reversed with an FDA approved drug that inhibits this receptor, a result with significant clinical implications. All together the findings reported here provide the first insights into the molecular and cellular mechanisms that close the temporal window for memory linking, thus segregating the memories for events that are temporally distinct.

## Supporting information

all figures

Virus vector information

Statistic information

## Methods

### Animals

*Ccr5* knockout (*Ccr5*^-/-^) mice were purchased from Taconic Farms (Germantown, NY). Experimental WT, *Ccr5*^+/-^ and *Ccr5*^-/-^ mice (3 to 5 months old) were generated by intercrossing *Ccr5*^+/-^ mice. Littermates were used for *Ccr5* KO linking test. 16-month-old male C57BL/6Nia were purchased from NIA for *Ccr5* expression detection and linking test. 11-week-old male C57BL/6N Tac mice were purchased from Taconic Farms (Germantown, NY) for all other experiments. Mice were group housed with free access to food and water, and maintained on a 12:12 hour light:dark cycle. All experiments were performed during the light phase of the cycle. All studies were approved by the Animal Research Committee at UCLA.

### Viral constructs

Constructs for *i*Tango2 system were gifts from Hyungbae Kwon, which include pAAV-hSYN-DRD2-V2tail-TevN-BLITz1-TetR-VP16-bGHpA (Addgene plasmid #89874; http://n2t.net/addgene:89874; RRID:Addgene_89874), pAAV-hSYN-bArrestin2-TevC-P2A-TdTomato-WPRE-bGHpA (Addgene plasmid #89873; http://n2t.net/addgene:89873; RRID:Addgene_89873), pAAV-TRE-EGFP (Addgene plasmid #89875; http://n2t.net/addgene:89875; RRID: Addgene_89875), pTRE-EGFP (Addgene plasmid #89871; http://n2t.net/addgene:89871; RRID: Addgene_89871). pGP-CMV-NES-jRGECO1a was a gift from Douglas Kim & GENIE Project (Addgene plasmid # 61563; http://n2t.net/addgene:61563; RRID: Addgene_61563). pAAV.Syn.GCaMP6f.WPRE.SV40 was a gift from Douglas Kim & GENIE Project (Addgene viral prep # 100837-AAV1; http://n2t.net/addgene:100837 ; RRID:Addgene_100837)

For the Opto-CCR5 experiment, pLenti-Ef1α-DIO-Opto-CCR5-EGFP was made by replacing the intracellular loops of rhodopsin with those of CCR5 to activate its specific intracellular signaling with light. The details of viral information are described in the Extended data Table 1.

### Real time-PCR

Total RNA was prepared using RNeasy Mini Kit (Qiagen, 74104) according to the manufacturer’s instructions. Single-stranded cDNA was synthesized using SuperScript III First-Strand Synthesis SuperMix (Invitrogen, 18080400). Real-time PCR was performed with SYBR Green-based reagents (iQ SYBR Green Supermix; Bio-Rad, 1708880) using a LightCycler 480 II (Roche). The following are primers used for real-time PCR:

Mouse *ccr5*, 5’GCTGCCTAAACCCTGTCATC-3’ and 5’GTTCTCCTGTGGATCGGGTA-3’

Mouse *ccl5*, 5′-TTCCACGCCAATTCATCGTT-3′ and 5′-GCATTCAGTTCCAGGTCAGTG-3′

Mouse *ccl3*, 5‘-AACCAAGTCTTCTCAGGGCC-3’ and 5’-CCAGGTCTCTTTGGAGTCAGC-3’

Mouse *ccl4*, 5’-CCAGGGTTCTCAGCACCAAT-3’ and 5’-TGGAGCAAAGACTGCTGGTC-3’

Mouse *ccl11*, 5’-AGATGCACCCTGAAA-3’ and 5’-GCATCCTGGACCCACT-3’

Mouse *36B4*, 5’-AGATCCAGCCAGATCCGCAT-3’ and 5’-GTTCTTGCCCATCAGCACC-3’

### In situ hybridization

Mouse brains were dissected and fast-frozen in OCT by dry Ice without PFA fixation. 20 μm frozen sections were sliced. In situ hybridization was performed using RNAscope Fluorescent Multiplex Reagent Kit (ACD, 320850) according to the manufacturer’s instructions. RNAscope Probe-Mm-Ccr5 (ACD, 438651) was used to detect *ccr5* mRNA. Probe-Mm-Rbfox3 (ACD, 313311) and Probe-Mm-Itgam-C2 (ACD, 311491) were used as markers for neurons and microglia, respectively.

### Immunostaining

Mice were transcardially perfused with 4% PFA (4% paraformaldehyte in 0.1 M phosphate buffer) and after perfusion, brains were sliced coronally (50 μm thick) with a vibratome and processed for immunostaining. Primary antibodies, including chicken polyclonal anti-GFP (Abcam AB13970, 1:1000), mouse anti-NeuN (Chemicon, MAB377, 1:1000), rabbit anti-GFAP (Dako, Z0334, 1:500), rabbit anti-cFos (Cell Signaling, 9F6, #2250, 1:500), and rabbit anti-P2Y12 (AnaSpec, AS-55043A, 1:1000) were used for immunostaining. Brain slices were incubated with 4’,6-diaminodino-2-phenylindole (DAPI, Invitrogen, 1:2000) for 10 min and washed with PBS three times before mounting onto slides. Immunostaining images were acquired with a Nikon A1 Laser Scanning Confocal Microscope (LSCM).

### Immunoblotting

Cultured HEK 293 cells were lysed with RIPA buffer (Sigma, St. Louis, MO, R0278) with protease inhibitor cocktail (Sigma, P8340), phosphatase inhibitor cocktail 2 (Sigma, P5726), phosphatase inhibitor cocktail 3 (Sigma, P0044). Protein samples (10 μg/well) were loaded to NuPAGE Novex 4–12% Bis-Tris protein gel (ThermoFisher Scientific, Carlsbad, CA, NP0336BOX) and transferred onto polyvinylidene difluoride (PVDF) membranes. The membranes were then blocked with 5% nonfat milk at room temperature for 1 hour and then probed with primary antibodies (phospho-p44/42 MAPK, Cell Signaling 9101, 1:4000, dilution) at 4°C overnight. Membranes were then incubated with HRP-conjugated secondary antibodies for 1 hour and developed with Supersignal solutions (Thermo Scientific). Then the membrane was stripped and probed again with primary antibodies (p44/42 MAPK, Cell Signaling 9102, 1:4000 dilution, β-actin 1:10,000, A5316, Sigma-Aldrich).

### CCR5-*i*Tango2 system

Inducible Tango (*i*Tango) system is a genetic method of labeling and manipulating cells with particular GPCR activation initially reported by Hyung-Bae Kwon lab^18^. Based on this method, we designed CCR5-*i*Tango2. Briefly, it couples a tetracycline-controlled transcriptional activator (tTA) to the C-terminal of mouse CCR5 via a specific tobacco etch virus (TEV) protease-sensitive cleavage site (TEV-seq), which is protected by AsLOV2/Jα (light sensitive domain). Upon activation, β-Arrestin tagged with TEV-C (C-terminal region of TEV) will bind intracellular loop of CCR5 tagged with TEV-N (N-terminal region of TEV), which will form functional TEV and cleave TEV-seq exposed to light stimulation. Then tTA will be released and translocate into nucleus to induce specific gene expression. To generate the CCR5-*i*Tango2 DNA constructs, full length mouse CCR5 cDNA was sub-cloned into pAAV-hSYN-DRD2-V2tail-TevN-BLITz1-TetR-VP16-bGHpA to replace DRD2 cDNA sequence (by VectorBuilder).

### Opto-CCR5 system

Opto-XR is the genetically encoded optical tool designed by Karl Deisseroth lab^19^, which can control GPCR-initiated biochemical signaling pathways with high spatiotemporal precision. Based on opto-XR, Won Do Heo lab designed and made the Opto-CCR5 construct and subclone it into a lentivirus backbone (Lenti-Ef1a-DIO-Opto-CCR5-EGFP). Briefly, the intracellular loops of rhodopsin were replaced with those of mouse CCR5. As a result, light induced structure change of rhodopsin would activate intracellular CCR5 signaling.

### Stereotaxic Surgery

Animals were anesthetized with 2% isoflurane and placed in a stereotaxic head frame on a heat pad. Artificial tears were applied to the eyes to prevent eye drying. A midline incision was made down the scalp, and a craniotomy was performed with a dental drill. After surgery, the animals were subcutaneously injected with Carprofen (5 mg/kg) and Dexamethasone (0.2 mg/kg) before recovery. Water with amoxicillin was applied for two weeks.

For cannula implantation, two guide cannulas (Plastics One, C313GS-5/SPC) were implanted at the following coordinates relative to bregma (mm): AP: −2.1, ML: ±1.7. Three weeks after cannulation, mice were anesthetized and sterilized PBS or mouse CCL5 peptide (70nM in PBS, 1 μL, 100nL/min) was infused into hippocampus through the internal cannula (Plastics One, C313IS-5/Spc) at DV:-1.6 relative to skull. After infusion, the internal cannula was left in place for an additional 5 min to ensure full diffusion.

For virus injection, a Nanoliter injector (World Precision Instruments) was used to infuse virus with Micro4 Controller (World Precision Instruments). Virus was infused at 50-100 nL/min. After infusion, the capillary was kept at the injection site for 5 min and then withdrawn slowly. The incision was closed with clips, which were removed 7 days later. The details of viruses used are described in the Supplemental Information (Table S1).

For optical fiber implantation, fiber Optic Cannula (Newdoon, 200 μm, NA=0.37) was immediately implanted after virus injection. The tip of the optic fiber was placed 600 μm above the virus injection site. Then, the canula was fixed with Metabond and dental cement.

For miniscope implantation, a GRIN lens was implanted into the dorsal hippocampal CA1 region as previously described^1^. After GCaMP6f virus injection, a ∼2mm diameter circular craniotomy was centered at the injection site. The cortex directly below the craniotomy was aspirated with a 27-gauge blunt syringe needle attached to a vacuum pump. Cortex buffer (NaCl 135mM, KCL 5mM, CaCl_2_ 2.5mM, MgSO_4_ 1.3mM, HEPES 5mM, PH 7.4) was repeatedly applied to the exposed tissue to prevent drying. The GRIN lens (0.50 NA, 2.0 mm in diameter, Grintech Gmbh) was slowly lowered above CA1 to a depth of 1.35 mm ventral to the surface of the skull at the most posterior point of the craniotomy. Next, a skull screw was used to anchor the lens to the skull. Both the lens and skull screw were fixed with super glue (Loctite, 45198) and dental cement (Jet Denture Repair Package, Lang, 1223CLR). Low Toxicity Silicone Adhesive (Kwik-Sil,World Precision Instruments) was used to cover the GRIN Lens for protection. Two weeks later, a small baseplate was cemented onto the animal’s head atop the previously formed dental cement.

### Optogenetics

For the CCR5-*i*Tango2 system, 3 weeks after virus injection and optic cannula implantation, the mice were handled for 3 days and then habituated with the optic fiber connected in their home cage for another 3 days (10min/day). Then the mice received contextual fear conditioning training and returned to their home cage. After 2.5h, 5.5h, 11.5h and 23.5h, different groups of mice received light stimulation in their home cage (473nm, 8-10mW, 10s on/50s off for 1h). The mice were kept for another 48h for GFP expression before the brains were collected and fixed with PFA perfusion.

For Opto-CCR5, the mice were anesthetized with 1.5% isoflurane during light delivery (473nm, ∼8mW, 50s on/10s off for 30min). Then, the mice were returned to their home cage for 30 min to recover before exposure to a different context.

### Behavioral procedures

The contextual memory linking task was carried out as previously described^1^. Mice were first handled for 3 days (1min/day) and then habituated to transportation and external environmental cues for 2 minutes in the experimental room each day for another 3 days. In the contextual memory linking task, mice explored 2 different contexts (A and then B) which were separated by 7 days or 5 hours. Mice explored each context for ten minutes, and the experiments with the 7-day and 5-hour intervals between contexts were counterbalanced. For immediate shock, mice were placed in chamber B for 10 s followed by a 2s shock (0.65 mA). 58 seconds after the shock, mice were placed back in their home cage. For the context tests, mice were returned to the designated context. Freezing was assessed via an automated scoring system (Med Associates) with 30 frames per second sampling; the mice needed to freeze continuously for at least one second before freezing could be counted.

### Slice preparation and CCL5 treatment

Adult mice (3-6 months old) were deeply anesthetized with isoflurane and the brains were rapidly dissected out and transferred to oxygenated (95% O2/5% CO2), ice-cold cutting solution containing 92 mM NaCl, 2.5 mM KCl, 1.25 mM NaH2PO4, 30 mM NaHCO3, 20 mM HEPES, 25 mM glucose, 2 mM Thiourea, 5m M Na-ascorbate, 3 mM Na-pyruvate, 2 mM CaCl2, and 2 mM MgCl2. Coronal slices (400 μm thick) containing the hippocampus were cut using a Leica VT1200 vibrating blade microtome, transferred to a submerged holding chamber containing oxygenated cutting solution and allowed to recover for 1h at room temperature. Prior to performing whole-cell recordings, each slice was incubated in a separate chamber containing either oxygenated aCSF (containing 115 mM NaCl, 10 mM glucose, 25.5 mM NaHCO3, 1.05 mM NaH2PO4, 3.3 mM KCl, 2 mM CaCl2, and 1 mM MgCl2) or 10nM CCL5 in oxygenated aCSF for 1h. Following incubation, slices were immediately transferred to a superfused recording chamber and constantly perfused with oxygenated aCSF maintained at 28°C. All recordings were performed within 30 min of aCSF or mouse CCL5 incubation.

### Whole-cell patch recordings

Whole cell current-clamp recordings were performed on pyramidal neurons in the CA1 region of the hippocampus using pipettes (3-5MΩ resistance) pulled from thin-walled Borosilicate glass using a Sutter P97 Flaming/Brown micropipette puller and filled with an internal solution containing 120 mM K-methylsuphate, 10 mM KCl, 10 mM HEPES, 10 mM Na-phosphocreatine, 4 mM Mg-ATP, and 0.4 mM Na-GTP. All recordings were obtained using a MultiClamp 700B amplifier controlled by the pClamp 10 software and digitized using the Digidata 1440A system. Signals were filtered at 10 kHz and digitized at 20 kHz. Neurons were included in the study only if the initial resting membrane potential (Vm) < -55 mV, access resistance (Ra) was < 20MΩ, and were rejected if the Ra changed by >20% of its initial value. For all recordings, neurons were held at -65 mV. The stable resting membrane potential of neurons was measured and averaged over a 60s duration with 0 mA current injection immediately after breaking in. To investigate the firing rate of neurons, the number of action potentials fired in response to a 600 msec pulse of depolarizing current injection (0 pA to 380 pA in 20 pA increments) was calculated. Three pulses were delivered for each current amplitude and the average number of action potentials fired for each current amplitude was plotted. The recordings were analyzed using Stimfit 0.15.8 and the data were screened for statistical outliers (± 2SD).

### Miniscope data acquisition and analyses

One-photon calcium imaging was recorded using UCLA miniscopes^33^. During recordings, digital imaging data were sent from the CMOS imaging sensor (Aptina, MT9V032) to custom data acquisition (DAQ) electronics and USB Host Controller (Cypress, CYUSB3013) over a light-weight, highly flexible co-axial cable. Images were acquired at 30 frames per second, using display resolution at 752 × 480 pixels (1 pixel = 1-2μm), and saved into uncompressed avi files. The analysis pipeline was written in MATLAB using first the NoRMCorre algorithm for motion correction (rigid registration)^34^, followed by individual neuron identification and extraction using the CNMF-E algorithm^35^. During motion correction, videos were 2x spatially down-sampled using the default built-in NoRMCorre protocol. During CNMF-E initialization, videos were further 2x spatially down-sampled and 5x temporally down-sampled. The quality of neuron extraction was verified using a MATLAB custom-made Neuron Deletion GUI. We excluded the detected putative neurons exhibiting ROI morphology or calcium trace abnormalities or incoherencies between the calcium trace peaks and the expected correspondent fluorescence increases in the video, and the neuron deletion was performed by experimenters blinded of the experimental groups and conditions. Each 10-min video from individual sessions was analyzed separately. Recordings from multiple sessions of the same animal were aligned using the spatial foot prints (neuron.A, output from CNMF-E) of each one of the detected cells for individual sessions. The centroid distance and spatial correlation were calculated for all cell pairs. Cell pairs from different sessions were considered to match if their spatial correlation ≤ 0.8 and their centroid distance ≤ 5 pixels. Overlapping percentages between two given sessions were calculated as the number of matched cells over the average of the total number of detected cells in each one of the two sessions. Overlapping Index= Ctx A^+^ Ctx B^+^ cell (Overlap) / [(Ctx A^+^ cell + Ctx B^+^ cell)/2] %.

### Quantification and Statistical Analyses

The investigators who collected and analyzed the data including behavior, electrophysiological and staining were blinded to the mouse genotypes and treatment conditions. Error bars in the figures indicate the SEM. All statistical analyses were performed using GraphPad Prism 6. For behavior and biochemical experiments, n designates the number of mouse or brains collected. For electrophysiological measurements, n designates the number of neurons. Statistical significance was assessed by Student’s t test, or one- or two-way ANOVA where appropriate, followed by the indicated post-hoc tests. The level of significance was set at p<0.05.

## Acknowledgements

We thank A. Macalino, E. Chen, E. Ramirez, C. Riviere-Cazaux, E. Lu, M. Tian for advice and technical support. This work was supported by grants from the NIMH (R01 MH113071), NIA (R01 AG013622), NINDS (RO1 NS106969) and from the Dr. Miriam and Sheldon G. Adelson Medical Research Foundation to A.J.S.

## Author contribution

YS and MZ did experimental design, data acquisition and analyses, drafting and revising the article; DC did memory linking time course and memory linking in aged *Ccr5* KO mice; GF did electrophysiology; YC and YS did qPCR; NK and WDH made the Opto-CCR5 construct; DN, CZ, AL, XK, and TS helped with data acquisition; DAF, AL and SH helped data analyses and interpretation; AJS did experimental design and interpretation, drafting and revising the article.

