## Supplementary material for "CCR5 closes the temporal window for memory linking": all figures

### Extended Data Fig. 1

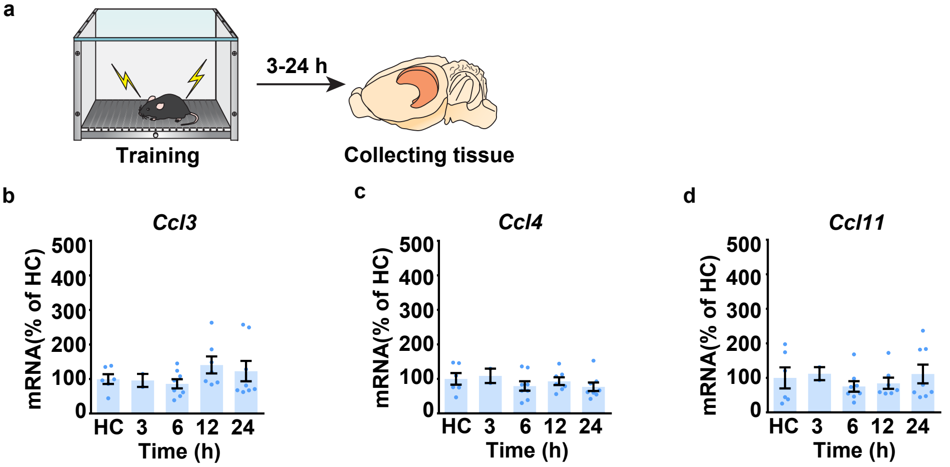


**Extended data Fig.1 Dorsal hippocampal *Ccl3*, *Ccl4*, *Ccl11* expression after fear conditioning.**

**a**, Schematics of hippocampal tissue collection.

**b**-**d**, qPCR experiment to measure *Ccl3* (**b**), *Ccl4* (**c**), and *Ccl11* (**d**) expression in naïve mice (HC) and in mice at different times after contextual fear conditioning. HC=home cage. n=2-8 mice per group.

Results show mean ± s.e.m.

### Extended Data Fig. 2

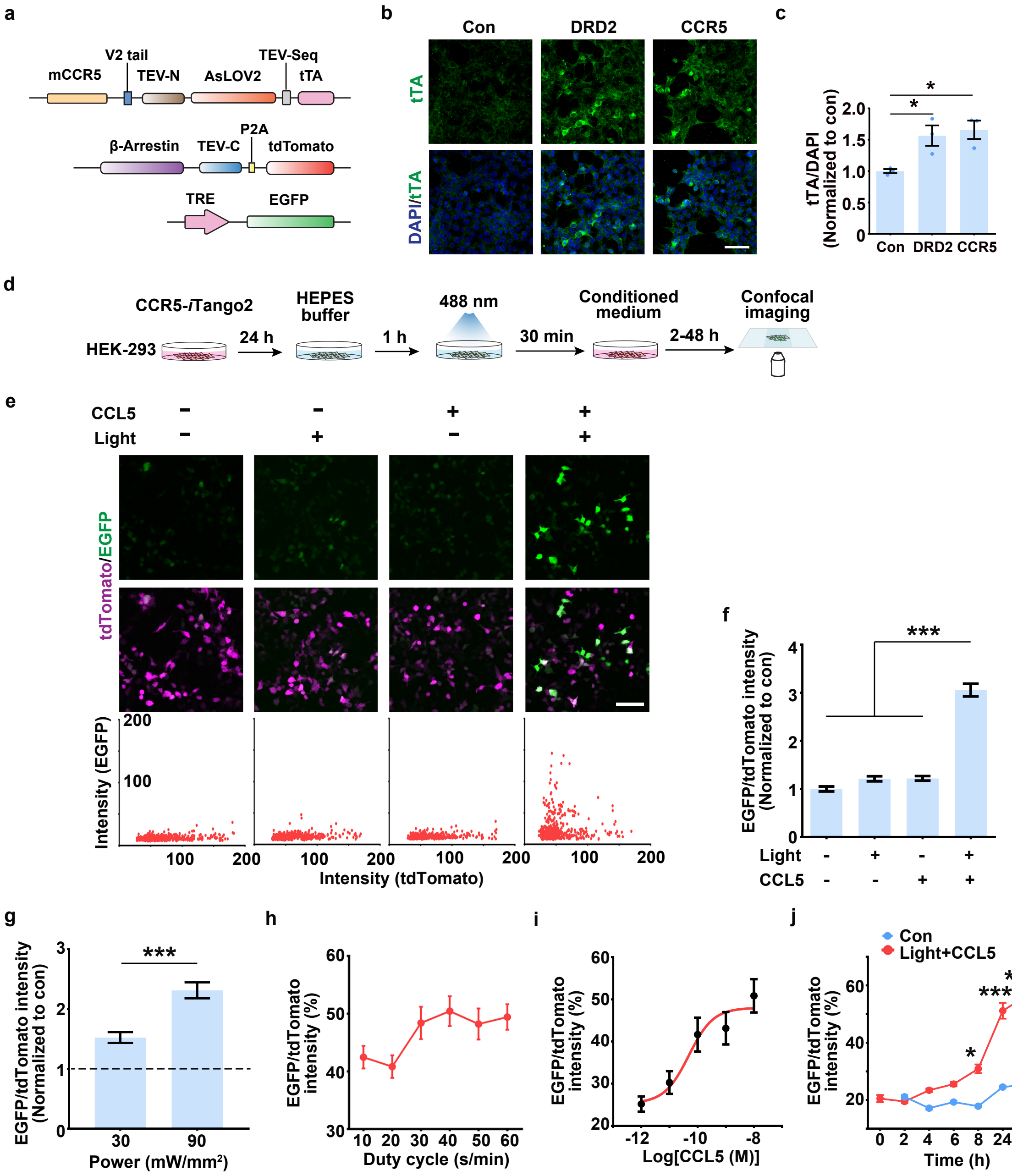

**Extended data Fig.2 Characterization of CCR5-*i*Tango2 *in vitro*.**

**a**, Schematics of CCR5-*i*Tango2 constructs.

**b, c**, Expression validation of the CCR5-*i*Tango system in HEK293 cells. DRD2-*i*Tango2 (for Dopamine 2 receptor) was used as a positive control. **b**, Representative images of tTA immunostaining. Scale bar, 50 μm. **c**, Quantification of tTA expression (intensity normalized to DAPI). n=3 slides per group, $^*P < 0.05$, one-way ANOVA.

**d**, HEK-293 cells were transfected with 3 plasmids (see methods) for 24h and then treated with 10 nM CCL5 and blue light to induce EGFP expression.

**e**, Representative images of EGFP expression after different treatments. Scale bar, 50 μm.

**f**, Quantification of EGFP and tdTomato ratio (intensity). n=70-282 cells per group, $^{***}P < 0.001$, one-way ANOVA. Compared to control, light or CCL5 group, only the group with both light and CCL5 showed EGFP expression.

**g**, Light power-dependent EGFP expression. Results were normalized to no light control. n=~300 cells per group. $^{***}P < 0.001$, student's t-test.

**h**, Duty cycle dependent EGFP expression. The light stimulation was delivered every minute (~0.017 Hz) to induce EGFP expression. Light was kept on for 10-60 s during each stimulation to induce EGFP expression.

**i**, Dose curve of CCL5 to induced CCR5 activation (measured by G/R fluorescence ratio) in cultured HEK293 cells.

**j**, Time course of EGFP expression. The green fluorescence increased monotonically during the different time intervals investigated. Compared to other time intervals (2, 4, 6, 8 and 24h), the 48h time interval showed the highest EGFP/tdTomato ratio. (n=~300 cells per group, $^*P<0.05$, $^{***}P<0.001$, two-way ANOVA).

Results show mean ± s.e.m.

#### Extended Data Fig. 3

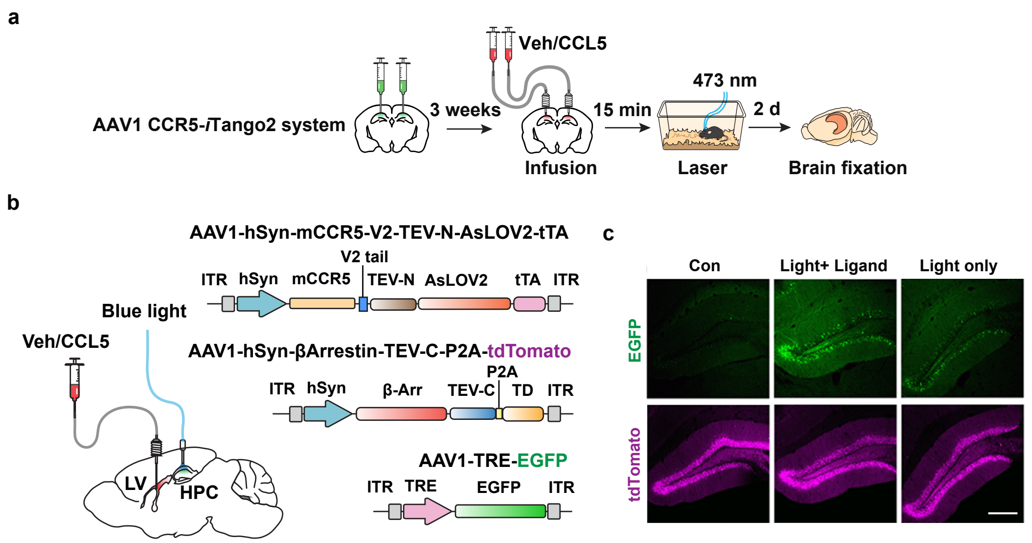


**Extended data Fig.3 CCR5-*i*Tango2 expression in vivo.**

**a**, Schematics of CCR5-*i*Tango2 AAVs injected into mouse hippocampus and validated through intra-hippocampal infusion of CCL5 and fiber-optic light delivery.

**b**, Left: To test CCR5-*i*Tango2 activation in dCA1 (Fig. 1h), CCL5 was infused into the lateral ventricle (LV) while light was delivered into dCA1 of hippocampus (HPC). Right: Schematics of CCR5-*i*Tango2 AAVs.

**c**, Representative images of CCR5-*i*Tango2-expressing hippocampal DG neurons in control condition (no light or CCL5), light only, and light with CCL5. Ligand and light were directly delivered into the hippocampus. Scale bar, 250 μm.

#### Extended Data Fig. 4

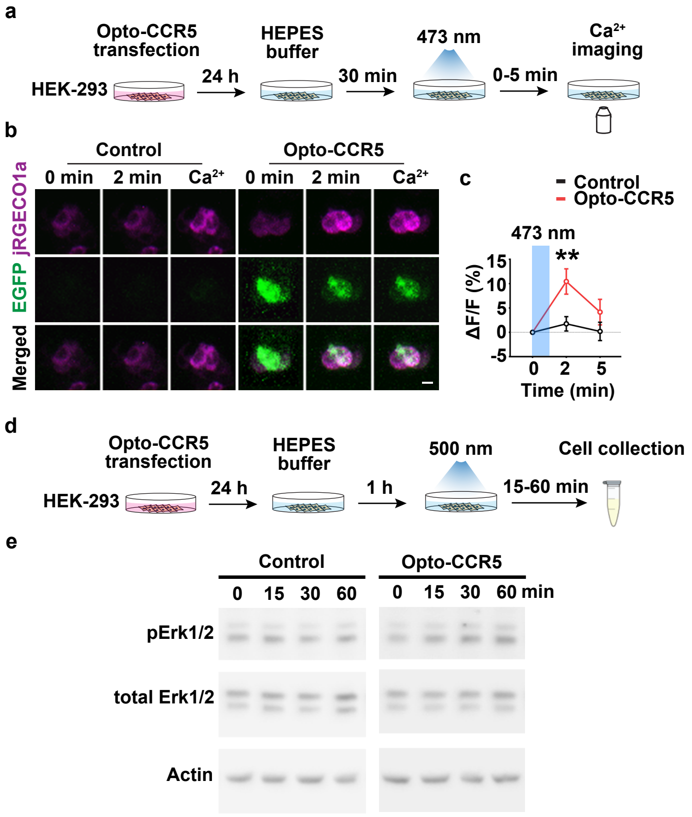


**Extended data Fig.4 Characterization of Opto-CCR5 *in vitro*.**

**a**, HEK-293 cells were transfected with Opto-CCR5 and jRGECO1a (Calcium sensor with red florescence) for 24h and then stimulated with blue light to induce a calcium response.

**b**, Representative images at 0 min or 2 min after stimulation, or in the medium with high calcium concentration. Scale bar, 20 μm.

**c**, Quantification of florescence change after light stimulation. In HEK-293 cells, Opto-CCR5-EGFP activation by light significantly increased intracellular $Ca^{2+}$ concentration reflected by jRGEC1a. n=~90 cells per group, $**P<0.01$, two-way ANOVA.

**d**, **e**, Opto-CCR5 activation increased pErk1/2 in HEK-293 cells.

**d**, HEK-293 cells were transfected with the Opto-CCR5 construct. After 24h expression, the cells were starved in HEPES buffer for 1h before a 2min light stimulation to reduce basal pErk1/2 levels.

**e**, Cells were collected at 0 (no light stimulation), 15, 30 or 60 min after light stimulation and subjected to Western blot analysis.

Results show mean ± s.e.m.

#### Extended Data Fig. 5

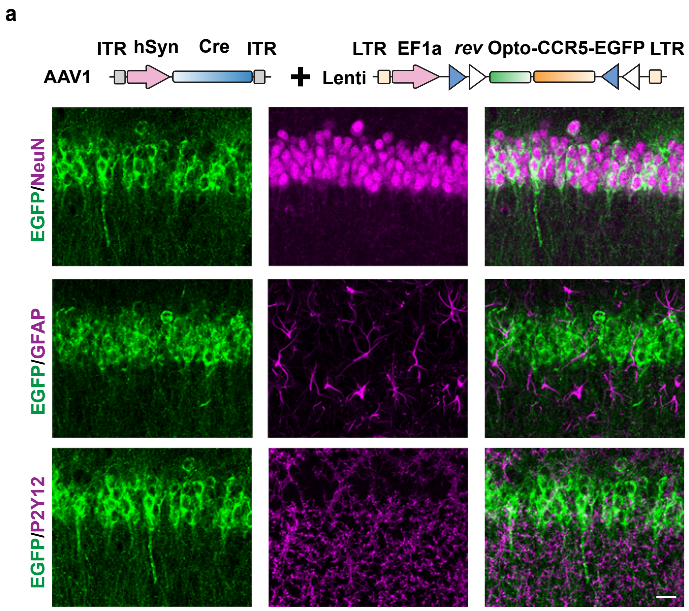


**Extended data Fig.5 Expression of Opto-CCR5 in dCA1 neurons.**
To limit Opto-CCR5 expression in dCA1 neurons, AAV1-hSyn-Cre was co-injected with Lenti-DIO-Opto-CCR5. NeuN (neuron marker), GFAP (astrocyte marker) and P2Y12 (microglia marker) were co-stained with EGFP in dCA1. Scale bar, 20 µm.

### Extended Data Fig. 6

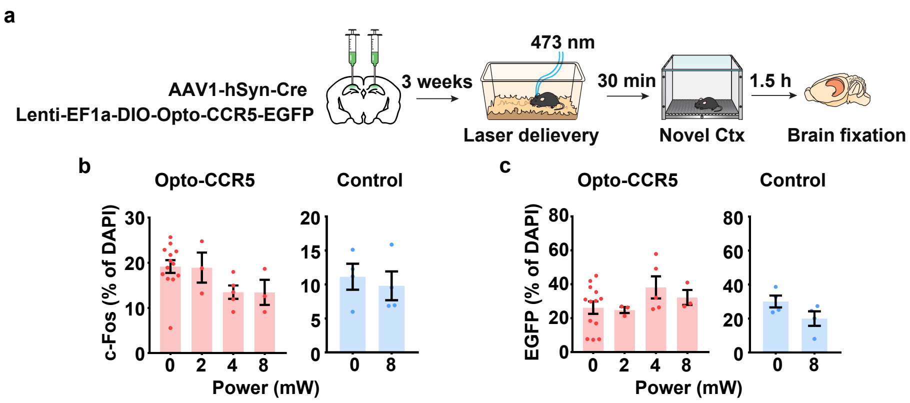


**Extended data Fig.6 Expression of c-Fos and Opto-CCR5 or EGFP control in dCA1.**
**a**, Percentage of c-Fos positive cells (normalized to cells with DAPI staining) in dCA1 with light stimulation of different power levels.
**b**, Percentage of EGFP expression cells (normalized to cells with DAPI staining) in dCA1with light stimulation of different power levels.
Results show mean ± s.e.m.

#### Extended Data Fig. 7

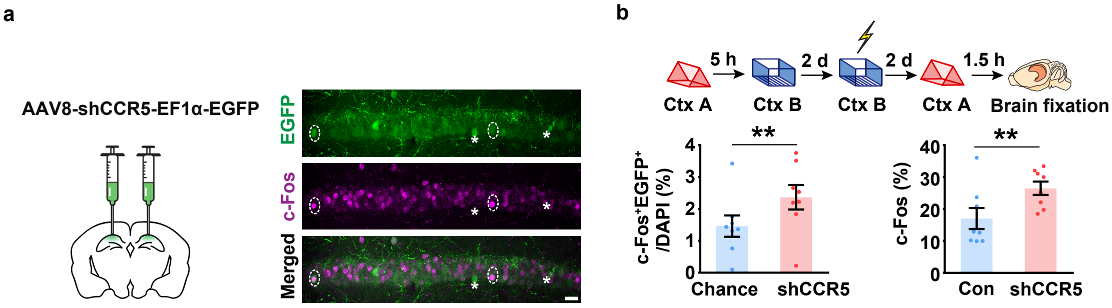


**Extended data Fig.7 *Ccr5* knockdown enhanced memory allocation.**

**a**, Schematics of AAV8-shRNA-CCR5-Ef1α-EGFP injection, and representative images of c-Fos and EGFP staining. Some $EGFP^{+}$c-$Fos^{+}$ were labelled by dotted line circle while some $EGFP^{+}$c-$Fos^{-}$ were labelled by asterisk. Scale bar, 20 μm.

**b**, dCA1 neurons with *Ccr5* knockdown had a higher probability of expressing c-Fos after a memory test in context A. Left: The percentage of c-$Fos^{+}EGFP^{+}$ cells in total (DAPI). Chance level was calculated as (c-$Fos^{+}$/DAPI)*($EGFP^{+}$/DAPI); right: percentage of c-$Fos^{+}$ cells in non-EGFP cells (Con) or in $EGFP^{+}$ cells with *Ccr5* knockdown (shCCR5). (n=8, $**P<0.01$, Student's t-test). Results show mean ± s.e.m.

Extended Data Fig. 8

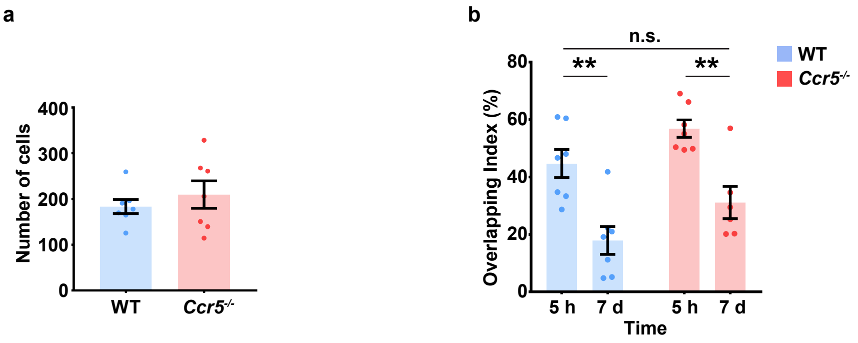


**Extended data Fig.8 Miniscope recording in WT and *Ccr5* KO mice.**

**a**, Total number of cells recorded in each WT and *Ccr5*$^{-/-}$ mouse

**b**, Overlapping index for WT and *Ccr5*$^{-/-}$ mice with a time interval of 5h or 7d (WT n=6, *Ccr5*$^{-/-}$ n=6; ***P*<0.001, two-way ANOVA).

All results shown as mean ± s.e.m.

### Extended Data Fig. 9

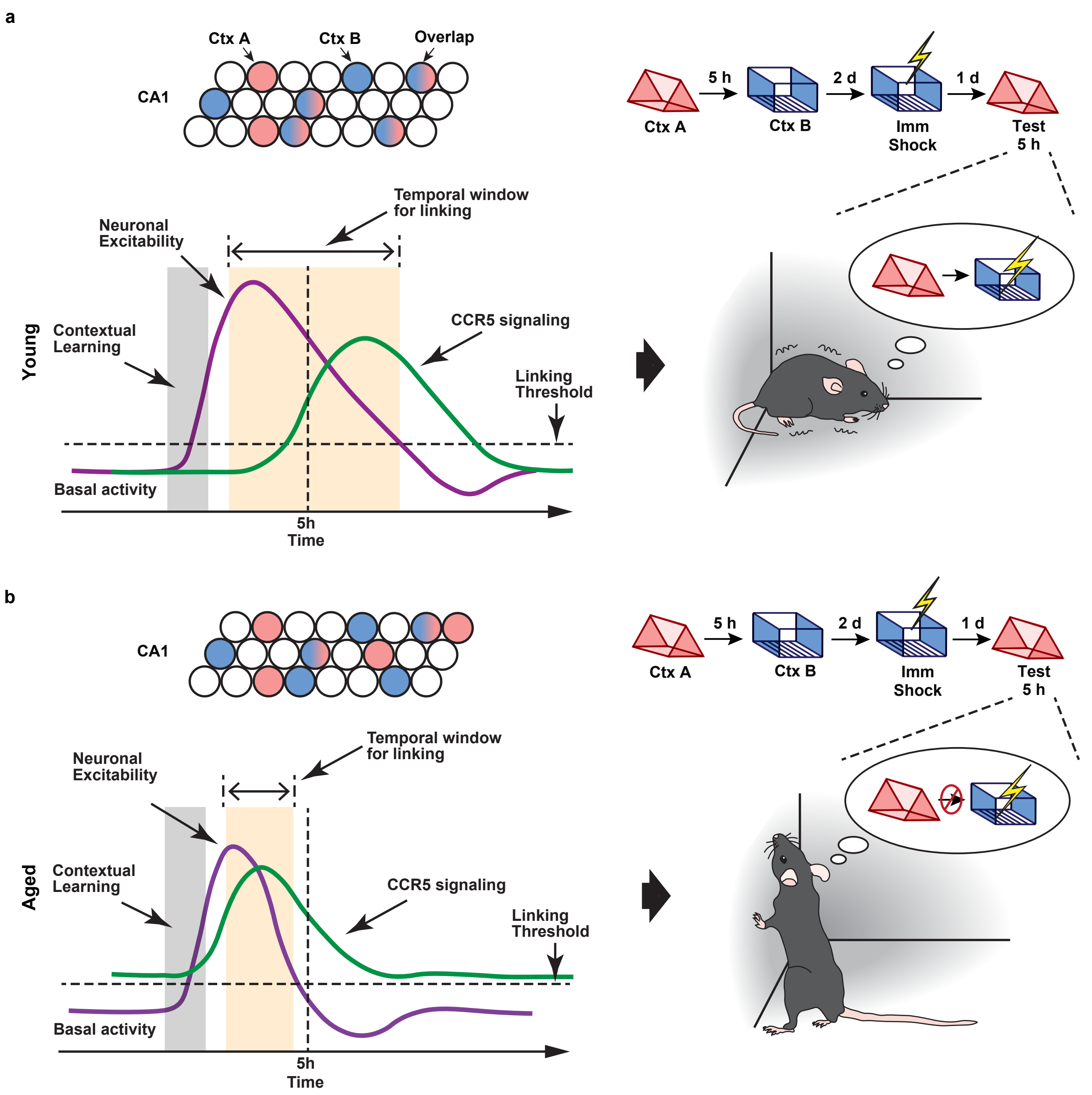


**Extended data Fig.9 Graphic abstract.**

**a**, In young mice, CCR5 signaling increases at a time point more than 5h after learning, and neuronal excitability and memory ensemble overlap remain high at 5h after learning. As a result, memories for context A (neutral context) and context B (shocked context) are linked together, and mice show high freezing during the test in context A.

**b**, In aged mice, CCR5 signaling is higher than young mice at baseline and there is a further increase before 5h after learning, which lead to a reduction of neuronal excitability and memory ensemble overlap at 5h after learning. As a result, memories for context A (neutral context) and context B (shocked context) are not linked, and mice show low freezing during the test in context A.
