## Supplementary material for "CCR5 closes the temporal window for memory linking": Virus vector information

### Supplemental Table 1, Stereotaxic coordinates for virus injection.

| Virus | Titer | Region |
|---|---|---|
| **CCR5-iTANGO** | | |
| *AAV1-hSYN-CCR5-V2tail-TevN-BLITz1-TetR-VP16-bGHpA* [a] | $9.5\times10^{12}$ TU/mL | Dorsal hippocampal CA1 |
| *AAV1-hSYN-bArrestin2-TevC-P2A-TdTomato-WPRE-bGHpA* [a] | $4.5\times10^{12}$ TU/mL | Dorsal hippocampal CA1 |
| *AAV1-TRE-EGFP* [a] | $4.7\times10^{11}$ TU/mL | Dorsal hippocampal CA1 |
| **OptoCCR5** | | |
| *Lenti-Ef1α-DIO-Opto-CCR5-EGFP* [b] | $5 \times 10^{8}$ TU/ml | Dorsal hippocampal CA1 |
| *AAV1-hSyn-CRE-WPRE-hGH* [c] | $1.0\times10^{11}$ TU/mL | |
| *AAV8-CAG-Flex-EGFP* [e] | $5.23\times10^{10}$ TU/mL | Dorsal hippocampal CA1 |
| *AAV1-hSyn-CRE-WPRE-hGH* [c] | $1.0\times10^{11}$ TU/mL | |
| **CCR5 Knock-down** | | |
| *AAV8-shRNA-CCR5-EGFP* [a] | $1.0\times10^{13}$ TU/mL | Dorsal hippocampal CA2/CA3* |
| *AAV8- shDsRed (shCon)-EGFP* [a] | $1.0\times10^{13}$ TU/mL | Dorsal hippocampal CA2/CA3* |
| **Miniscope** | | |
| *AAV1-Syn-GCaMP6f-WPRE-SV40* [d] | $1.7\times10^{13}$ TU/mL | Dorsal hippocampal CA1 |

[a]The virus was prepared by Yang Shen according to protocol from Dr. Corinna Burger (University of Wisconsin-Ma

| Coordinates | Volume |
|---|---|
| AP: −2.1 mm, ML: ±2.0 mm relative to bregma; DV: −1.65 mm relative to skull | 250 nL per point |
|  | 250 nL per point |
|  | 500 nL per point |
| AP: −2.1 mm, ML: ±2.0 mm relative to bregma; DV: −1.65 mm relative to skull | 950 nL per point |
|  | 50 nL per point |
| AP: −2.1 mm, ML: ±2.0 mm relative to bregma; DV: −1.65 mm relative to skull | 950 nL per point |
|  | 50 nL per point |
| AP: −1.8 mm, ML: ±1.4 mm relative to bregma; DV: −1.65 mm relative to skull; AP: −2.5 mm, ML: ±2.0 mm relative to bregma; DV: −1.65 mm relative to skull | 1.5 μL per point |
| AP: −2.1 mm, ML: −2.0 mm relative to bregma; DV: −1.65 mm relative to skull | 1 μL |

dison); [b]The virus was pepared by Masakazu Kamata (University of Alabama at Birmingham); [c]Upenn vector Core;

[d]Addgene; [e]Salk vector Core
