## Supplementary material for "CCR5 closes the temporal window for memory linking": Statistic information

| Figure | Panel | Test | Group size | Statistic | P value | Pair-wise | Statistic2 |
|---|---|---|---|---|---|---|---|
| **Fig, 1** | **b** | One-way ANOVA | HC: 18<br>3 h: 7<br>6 h: 8<br>12 h: 8<br>24 h: 10 | F(4, 46) = 3.133 | 0.0232 | Dunnett's multiple comparisons test | HC vs. 3 h q(46)=0.04135, p>0.9999<br>HC vs. 6 h q(46)=1.253, p=0.5364<br>HC vs. 12 h q(46)=2.798, p=0.0259<br>HC vs. 24 h q(46)=2.571, p=0.0453 |
| | **c** | One-way ANOVA | HC: 11<br>3 h: 4<br>6 h: 8<br>12 h: 8<br>24 h: 8 | F(4, 34) = 2.759 | 0.0434 | Dunnett's multiple comparisons test | HC vs. 3 h q(34)=0.1588, p=0.9994<br>HC vs. 6 h q(34)=2.022, p=0.1558<br>HC vs. 12 h q(34)=2.61, p=0.0448<br>HC vs. 24 h q(34)=0.1838, p=0.999 |
| | **e** | Unpaired t test (Two -tailed) | Neuron: 5<br>Microglia: 5 | t(8) =2.355 | 0.0463 | N/A | |
| | **f** | 2-way ANOVA (Time X Cell type) | Neuron HC: 5<br>Neuron 3 h: 4<br>Neuron 6 h: 5<br>Neuron 12 h: 4<br>Neuron 24 h: 4 | Main effect of time<br>F(4, 34) = 3.675 | 0.0136 | N/A | |
| | | | Microglia HC: 5<br>Microglia 3 h: 4<br>Microglia 6 h: 5<br>Microglia 12 h: 4<br>Microhglia 24 h:4 | Main effect of cell type F(1, 34) = 1.404 | 0.2443 | N/A | |
| | | | | Time X Cell type<br>F(4, 34) = 2.194 | 0.0905 | Dunnett's multiple comparisons test | Neuron HC vs. 3 h q(34)=0.0491, p>0.9999<br>Neuron HC vs. 6 h q(34)=3.432, p=0.0058<br>Neuron HC vs. 12 h q(34)= 3.038 p=0.016<br>Neuron HC vs. 24 h q(34)=0.132, p=0.9998<br>Microglia HC vs. 3 h q(34)=0.09693, p=0.9999<br>Microglia HC vs. 6 h q(34)=0.2002, p=0.9988<br>Microglia HC vs. 12 h q(34)= 0.5009, p=0.9636<br>Microglia HC vs. 24 h q(34)=0.1767, p=0.9991 |
| | **j** | One-way ANOVA | HC: 5<br>3 h: 6<br>6 h: 6<br>12 h: 5<br>24 h: 5 | F(4, 22) = 2.627 | 0.0621 | Dunnett's multiple comparisons test | HC vs. 3 h q(22)=1.234, p=0.556<br>HC vs. 6 h q(22)=1.955, p=0.1872<br>HC vs. 12 h q(22)=2.215, p=0.1156<br>HC vs. 24 h q(22)=3.025, p=0.0212 |
| **Fig. 2** | **a** | One-way ANOVA | 5 h: 32<br>1 d: 26<br>2 d: 14<br>7 d: 16 | F(3, 84) = 3.957 | 0.0109 | Dunnett's multiple comparisons test | 5 h vs. 1 d q(84)=2.412, p=0.0477<br>5 h vs. 2 d q(84)=2.745, p=0.0202<br>5 h vs. 7 d q(84)=2.609, p=0.0290 |
| | **b** | Two-way repeated measures ANOVA (Treatment X Context, matching across treatment) | Veh: 20<br>CCL5 :17 | Main effect of treatment<br>F(1, 35) = 4.207 | 0.0478 | N/A | |
| | | | | Main effect of context<br>F(2, 70) =56.07 | <0.0001 | N/A | |
| | | | | Interaction<br>F(2, 70) = 1.252 | 0.2922 | Sidak's multiple comparisons test | Veh Shocked vs. Novel t(70)=8.081, p<0.0001<br>Veh 5 h vs. Novel t(70)=5.294, p<0.0001<br>CCL5 Shocked vs. Novel t(70)=6.941, p<0.0001<br>CCL5 5 h vs. Novel t(70)=2.815, p=0.0126 |
| | | | | Subjects (matching)<br>F(35, 70) = 1.966 | 0.0082 | Sidak's multiple comparisons test | Shocked Veh vs CCL5 t(105)=1.128, p=0.5978<br>5 h Veh vs CCL5 t(105)=2.536, p=0.0376<br>Novel Veh vs CCL5 t(105)=0.6682, p=0.8791 |
| | **e** | Two-way repeated measures ANOVA (Treatment X Context,matching across treatment) | Control: 15<br>Opto-CCR5: 14 | Main effect of treatment<br>F(1, 27) = 0.03677 | 0.8494 | N/A | |
| | | | | Main effect of context<br>F(2, 54) =21.75 | <0.0001 | N/A | |
| | | | | Interaction<br>F(2, 54) = 0.9364 | 0.3983 | N/A | |
| | | | | Subjects (matching)<br>F(27, 54) = 5.719 | 0.0001 | Sidak's multiple comparisons test | shCon Shocked vs. Novel t(54)=4.085, p=0.0004<br>shCon 5 h vs. Novel t(54)=0.8135, p=0.8044<br>shCon 5 h vs. Shocked t(54)=3.271, p=0.0056<br>shCCR5 Shocked vs. Novel t(54)=5.242, p<0.0001<br>shCCR5 5 h vs. Novel t(54)=2.917, p=0.0153<br>shCCR5 5 h vs. Shocked t(54)=2.325, p=0.07 |
| | **g** | Two-way repeated measures ANOVA (Treatment X Context, matching across treatment) | shCon: 13<br>shCCR5 :16 | Main effect of treatment<br>F(1, 35) = 4.207 | 0.0478 | N/A | |
| | | | | Main effect of context<br>F(2, 70) =56.07 | <0.0001 | N/A | |

| | | | | | | | |
|---|---|---|---|---|---|---|---|
| | | | | Interaction<br>F(2, 70) = 1.252 | 0.2922 | Sidak's multiple comparisons test | Veh Shocked vs. Novel t(70)=8.081, p<0.0001<br>Veh 5 h vs. Novel t(70)=5.294, p<0.0001<br>CCL5 Shocked vs. Novel t(70)=6.941, p<0.0001<br>CCL5 5 h vs. Novel t(70)=2.815, p=0.0126 |
| | | | | Subjects (matching)<br>F(35, 70) = 1.966 | 0.0082 | Sidak's multiple comparisons test | Shocked Veh vs CCL5 t(105)=1.128, p=0.5978<br>5 h Veh vs CCL5 t(105)=2.536, p=0.0376<br>Novel Veh vs CCL5 t(105)=0.6682, p=0.8791 |
| | **h** | Two-way repeated measures ANOVA (Genotype X Context,matching across genotype) | WT: 9<br>*Ccr5+/-* : 6<br>Ccr5-/- : 7 | Main effect of genotype<br>F(2, 19) = 0.8974 | 0.4242 | N/A | |
| | | | | Main effect of context<br>F(2, 38) =5.106 | 0.0109 | N/A | |
| | | | | Interaction<br>F(4, 38) = 0.9093 | 0.4684 | N/A | |
| | | | | Subjects (matching)<br>F(19, 38) = 10.49 | <0.0001 | Holm-Sidak's multiple comparisons test | WT Shocked vs. 7 d(38)=3.444, p=0.0028<br>WT 5 h vs. 7 d t(38)=2.153, p=0.0378<br>Ccr5+/- Shocked vs. 7 d t(38)=1.431, p=0.2607<br>Ccr5+/- 5 h vs. 7 d t(38)=1.507, p=0.2607<br>Ccr5-/- Shocked vs. Novel t(38)=0.6277, p=0.7828<br>Ccr5-/- 5 h vs. 7 d t(38)=0.5579, p=0.7828 |
| **Figure 3** | **b** | Two-way repeated measures ANOVA (Treatment X Current injection, matching across treatment) | Veh: 10<br>CCL5 :9 | Main effect of treatment<br>F(1, 17) = 4.487 | 0.0492 | N/A | |
| | | | | Main effect of current injection<br>F(4, 68) =3.113 | 0.0206 | N/A | |
| | | | | Interaction<br>F(4, 68) = 0.8686 | 0.2922 | N/A | |
| | | | | Subjects (matching)<br>F(17, 68) = 323.8 | <0.0001 | N/A | |
| | **d** | Two-way repeated measures ANOVA (treatment X Power,matching across Power) | 0 mW: 13<br>2 mW: 3<br>4 mW: 5<br>8 mW: 3 | Main effect of treatment<br>F(1, 20) = 53.74 | <0.0001 | N/A | |
| | | | | Main effect of Power<br>F(3, 20) = 0.3555 | 0.7857 | N/A | |
| | | | | Interaction<br>F(3, 20) = 0.5098 | 0.68 | N/A | |
| | | | | Treatment X Genotype: F(1, 27) = 13.07 | 0.0012 | N/A | |
| | **e** | One-way ANOVA (Opto-CCR5) | 0 mW: 13<br>2 mW: 3<br>4 mW: 5<br>8 mW: 3 | F(3, 19) = 15.21 | <0.0001 | Dunnett's multiple comparisons test | 0 mW vs. 2 mW q(19)=0.09477, p=0.9994<br>0 mW vs. 4 mW q(19)=5.509, p<0.0001<br>0 mW vs. 8 mW q(19)=4.694, p=0.0005 |
| | | Unpaired t test (EGFP, Two -tailed) | 0 mW: 4<br>8 mW: 4 | t(6) =0.4207 | 0.6886 | N/A | |
| | **f** | One-way ANOVA (Opto-CCR5) | 0 mW: 13<br>2 mW: 3<br>4 mW: 5<br>8 mW: 3 | F(3, 19) = 9.395 | 0.0005 | Dunnett's multiple comparisons test | 0 mW vs. 2 mW q(19)=0.09796, p=0.9993<br>0 mW vs. 4 mW q(19)=3.988, p=0.0022<br>0 mW vs. 8 mW q(19)=3.999, p=0.0021 |
| | | Unpaired t test (EGFP, Two -tailed) | 0 mW: 4<br>8 mW: 4 | t(6) =0.8635 | 0.421 | N/A | |
| | **i** | 2-way ANOVA (Genotype X Time) | WT 5 h: 6<br>WT 1 d: 6<br>WT 2 d: 6<br>WT 7 d: 6 | Main effect of genotype F(3, 45) = 10.52 | <0.0001 | N/A | |
| | | | Ccr5-/- 5 h: 6<br>Ccr5-/- 1 d: 6<br>Ccr5-/- 2 d: 6<br>Ccr5-/- 7 d: 6 | Main effect of time<br>F(1, 45) = 10.73 | 0.002 | N/A | |
| | | | | Genotype X Time<br>F(3, 45) = 0.05513 | 0.9827 | N/A | |
| **Figure 4** | **a** | Unpaired t test (*Ccr5* , Two -tailed) | Young: 14<br>Aged: 6 | t(18) =2.175 | 0.0432 | N/A | |
| | | Unpaired t test (*Ccl5* , Two -tailed) | 0 mW: 12<br>8 mW: 5 | t(15) =9.272 | <0.0001 | N/A | |
| | **b** | One-way ANOVA (*Ccr5* ) | HC: 6<br>3 h: 6<br>6 h: 6 | F(2, 15) = 0.0711 | 0.9317 | Tukey's multiple comparisons test | HC vs. 3 h q(15)=0.4958, p=0.9348<br>HC vs. 6 h q(15)=0.418, p=0.9531<br>3 h vs. 6 h q(15)=0.07787, p=0.9983 |

| | | | | | | | |
|---|---|---|---|---|---|---|---|
| | | One-way ANOVA<br>(*Ccl5* ) | HC: 5<br>3 h: 6<br>6 h: 6 | F(2, 14) = 8.598 | 0.0037 | Tukey's multiple comparisons test | HC vs. 3 h q(14)=4.736, p=0.0124<br>HC vs. 6 h q(14)=0.3155, p=0.973<br>3 h vs. 6 h q(14)=5.298, p=0.0058 |
| | **c** | Two-way repeated measures ANOVA (Genotype X Context, matching across Genotype) | WT: 7<br>*Ccr5-/-* : 8 | Main effect of genotype<br>F(1, 13) = 0.06446 | 0.8035 | N/A | |
| | | | | Main effect of context<br>F(2, 26) =23.9 | <0.0001 | N/A | |
| | | | | Interaction<br>F(2, 26) = 2.768 | 0.0813 | N/A | |
| | | | | Subjects (matching)<br>F(13, 26) = 5.318 | 0.0001 | Sidak's multiple comparisons test | WT Shocked vs. 5 h t(26)=4.275, p=0.0007<br>WT Shocked vs. Novel t(26)=4.524, p=0.0004<br>WT 5 h vs. Novel t(26)=0.2498, p=0.9926<br>*Ccr5-/-* Shocked vs. 5 h(26)=1.766, p=0.2445<br>*Ccr5-/-* Shocked vs. Novel t(26)=5.167, p<0.0001<br>*Ccr5-/-* 5 h vs. Novel t(26)=3.401, p=0.0065 |
| | **d** | Two-way repeated measures ANOVA (Treatment X Context, matching across treatment) | Veh: 15<br>Maraviroc: 14 | Main effect of treatment<br>F(1, 27) = 3.085 | 0.0904 | N/A | |
| | | | | Main effect of context<br>F(2, 54) = 9.54 | 0.0003 | N/A | |
| | | | | Interaction<br>F(2, 54) = 2.354 | 0.1047 | Tukey's multiple comparisons test | Veh Shocked vs. 5 h q(54)=4.576, p=0.0058<br>Veh Shocked vs. Novel q(54)=4.053, p=0.016<br>Veh 5 h vs. Novel q(54)=0.523, p=0.9275<br>Maraviroc Shocked vs. 5 h q(54)=1.083, p=0.7256<br>Maraviroc Shocked vs. Novel q(54)=4.547, p=0.0061<br>Maraviroc 5 h vs. Novel q(54)=3.464, p=0.0455 |
| | | | | Subjects (matching)<br>F(27, 54) = 3.839 | <0.0001 | Sidak's multiple comparisons test | Shocked Veh vs Maraviroc t(81)=0.959, p=0.713<br>5 h Veh vs Maraviroc t(81)=2.68, p=0.0265<br>Novel Veh vs Maraviroc t(81)=0.6334, p=0.895 |
| **Extended fig. 1** | **a** | One-way ANOVA | HC: 6<br>3 h: 2<br>6 h: 8<br>12 h: 7<br>24 h: 8 | F(4, 26) = 0.9655 | 0.4431 | Dunnett's multiple comparisons test | HC vs. 3 h q(26)=0.08098, p=0.9999<br>HC vs. 6 h q(26)=0.425, p=0.9794<br>HC vs. 12 h q(26)=1.247, p=0.5454<br>HC vs. 24 h q(26)=0.7272, p=0.8779 |
| | **b** | One-way ANOVA | HC: 6<br>3 h: 2<br>6 h: 8<br>12 h: 7<br>24 h: 8 | F(4, 26) = 0.6653 | 0.6218 | Dunnett's multiple comparisons test | HC vs. 3 h q(26)=0.3011, p=0.9944<br>HC vs. 6 h q(26)=1.053, p=0.6782<br>HC vs. 12 h q(26)=0.3297, p=0.9919<br>HC vs. 24 h q(26)=1.174, p=0.595 |
| | **c** | One-way ANOVA | HC: 6<br>3 h: 2<br>6 h: 8<br>12 h: 7<br>24 h: 8 | F(4, 26) = 0.4676 | 0.7589 | Dunnett's multiple comparisons test | HC vs. 3 h q(26)=0.2503, p=0.9971<br>HC vs. 6 h q(26)=0.7651, p=0.8583<br>HC vs. 12 h q(26)=0.463, p=0.9721<br>HC vs. 24 h q(26)=0.3495, p=0.9899 |
| **Extended fig. 2** | **c** | One-way ANOVA | Con: 3<br>DRD2: 3<br>CCR5: 8 | F(2, 6) = 7.818 | 0.0213 | Dunnett's multiple comparisons test | Con vs. CCR5 q(6)=3.652, p=0.0189<br>Con vs. DRD2 q(6)=3.138, p=0.0353 |
| | **f** | One-way ANOVA | Light-CCL5-: 70<br>Light+CCL5-: 97<br>Light-CCL5+: 97<br>Light+CCL5+: 282 | F(3, 542) = 61.86 | <0.0001 | Dunnett's multiple comparisons test | Light-CCL5- vs. Light+CCL5+ q(542)=9.443, p<0.0001<br>Light+CCL5- vs. Light+CCL5+ q(542)=9.61, p<0.0001<br>Light-CCL5+ vs. Light+CCL5+ q(542)=9.568, p<0.0001 |
| | **g** | Unpaired t test (Two -tailed) | 30: 320<br>90: 307 | t(625) =4.939 | <0.0001 | N/A | |
| | **j** | 2-way ANOVA (Time X Treatment) | Light+CCL5+ 0 h: 58<br>Light+CCL5+ 2 h: 194<br>Light+CCL5+ 4 h: 282<br>Light+CCL5+ 6 h: 310<br>Light+CCL5+ 8 h: 316<br>Light+CCL5+ 24 h: 396<br>Light+CCL5+ 48 h: 345 | Main effect of time<br>F(6, 3799) = 39.69 | <0.0001 | N/A | |

| | | | | | | | |
|---|---|---|---|---|---|---|---|
| | | | Light-CCL5- 2 h: 195<br>Light-CCL5- 4 h: 219<br>Light-CCL5- 6 h: 290<br>Light-CCL5- 8 h: 304<br>Light-CCL5- 24 h: 445<br>Light-CCL5- 48 h: 401 | Main effect of TREATMENT<br>F(1, 3799) = 123.5 | <0.0001 | N/A | |
| | | | | Time X Cell type<br>F(6, 3799) = 20.62 | <0.0001 | Sidak's multiple comparisons test | Light+CCL5+ 0 h vs. 2 h t(3799)=0.24515, p>0.9999<br>Light+CCL5+ 0 h vs. 4 h t(3799)=0.7497, p=0.9733<br>Light+CCL5+ 0 h vs. 6 h t(3799)=1.312, p=0.717<br>Light+CCL5+ 0 h vs. 8 h t(3799)=2.7, p=0.041<br>Light+CCL5+ 0 h vs. 24 h t(3799)=8.086, p<0.0001<br>Light+CCL5+ 0 h vs. 48 h t(3799)=9.414, p<0.0001 |
| **Extended fig. 4** | **c** | 2-way ANOVA (Treatment X Time) | Control 2 min: 95<br>Control 5 min: 96<br>Opto-CCR5 2 min: 86<br>Opto-CCR5 2 min: 89 | Main effect of treatment<br>F(1, 362) = 8.524 | 0.0037 | N/A | |
| | | | | Main effect of time<br>F(1, 362) = 3.281 | 0.0709 | N/A | |
| | | | | Treatment X Time<br>F(1, 362) = 1.19 | 0.276 | Sidak's multiple comparisons test | 2 min Control vs. Opto-CCR5 t(362)=2.82, p=0.0101<br>5 min Control vs. Opto-CCR5 t(362)=1.301, p=0.3507 |
| **Extended fig. 6** | **b** | One-way ANOVA (Opto-CCR5) | 0 mW: 13<br>2 mW: 3<br>4 mW: 5<br>8 mW: 3 | F(3, 20) = 2.424 | 0.0958 | Dunnett's multiple comparisons test | 0 mW vs. 2 mW q(20)=0.07642, p=0.9996<br>0 mW vs. 4 mW q(20)=2.229, p=0.0931<br>0 mW vs. 8 mW q(20)=1.846, p=0.1883 |
| | | Unpaired t test (EGFP, Two -tailed) | 0 mW: 4<br>8 mW: 4 | t(6) =0.4681 | 0.6562 | N/A | |
| | **c** | One-way ANOVA (Opto-CCR5) | 0 mW: 13<br>2 mW: 3<br>4 mW: 5<br>8 mW: 3 | F(3, 20) = 1.296 | 0.3031 | Dunnett's multiple comparisons test | 0 mW vs. 2 mW q(20)=0.177, p=0.996<br>0 mW vs. 4 mW q(20)=1.824, p=0.1954<br>0 mW vs. 8 mW q(20)=0.7514, p=0.7949 |
| | | Unpaired t test (EGFP, Two -tailed) | 0 mW: 4<br>8 mW: 4 | t(6) =1.82 | 0.1185 | N/A | |
| **Extended fig. 7** | **b** | Paired t test (Left, Two -tailed) | n= 8 | t(7) =3.874 | 0.0061 | N/A | |
| | | Paired t test (Right, Two -tailed) | n= 8 | t(7) =3.974 | 0.0054 | N/A | |
| **Extended fig. 8** | **a** | Unpaired t test (Two -tailed) | WT: 7<br>Ccr5-/-: 7 | t(12) = 0.7788 | 0.4512 | N/A | |
| | **b** | 2-way ANOVA (Genotype X Time) | WT 5h: 7<br>WT 7d: 7<br>*Ccr5-/-* 5h: 7<br>Ccr5-/- 7d: 6 | Main effect of Genotype F(1, 23) =32 | <0.0001 | N/A | |
| | | | | Main effect of time<br>F(1, 23) = 7.485 | 0.0118 | N/A | |
| | | | | Genotype X Time<br>F(1, 23) = 0.0122 | 0.913 | Sidak's multiple comparisons test | WT 5h vs. *Ccr5-/-* 5h t(23)=1.895, p=0.3562<br>WT 5h vs. WT 7d t(23)=4.162, p=0.0023<br>WT 5h vs. *Ccr5-/-* 7d t(23)=2.026, p=0.2859<br>*Ccr5-/-* 5h vs. Wt 7d t(23)=6.057, p=<0.0001<br>*Ccr5-/-* 5h vs. *Ccr5-/-* 7d t(23)=3.846, p=0.0049<br>WT 7d vs. *Ccr5-/-* 7d t(23)=1.974, p=0.3126 |
